# GDF15 antagonism limits severe heart failure and prevents cardiac cachexia in mice

**DOI:** 10.1101/2022.09.06.506633

**Authors:** Minoru Takaoka, John A. Tadross, Ali Al-Hadithi, Rocío Villena-Gutiérrez, Jasper Tromp, Shazia Absar, Marcus Au, James Harrison, P. Coll Anthony, Stefan J. Marciniak, Debra Rimmington, Eduardo Oliver, Borja Ibáñez, A. Voors Adriaan, Stephen O’Rahilly, Ziad Mallat, Jane C. Goodall

**Affiliations:** Heart and Lung Research Institute. Department of Medicine, University of Cambridge, Cambridge, UK; Wellcome-MRC Institute of Metabolic Science and Medical Research Council, Metabolic Diseases Unit, University of Cambridge, Cambridge, UK; Department of Histopathology and East Midlands & East of England Genomic Laboratory; Centro Nacional de Investigaciones Cardiovasculares (CNIC), Madrid. Spain; University of Groningen, University Medical centre Groningen, the Netherlands; National Heart Centre Singapore, Singapore; Cambridge Institute for Medical Research, Cambridge Biomedical Campus, University of Cambridge, Cambridge, UK; Centro de Investigaciones Biomédicas en Red de Enfermedades Cardiovasculares (CIBERCV), Madrid. Spain; Centro de Investigaciones Biologicas Margarita Salas (CIB-CSIC), Madrid, Spain; IIS-Hospital Fundacion Jimenez Diaz, Madrid, Spain

## Abstract

Heart failure and associated cachexia is an unresolved and important problem. We report a new model of severe heart failure that consistently results in cachexia. Mice lacking the integrated stress response (ISR) induced eIF2α phosphatase, PPP1R15A, exhibit a dilated cardiomyopathy and severe weight loss following irradiation, whilst wildtype mice are unaffected. This is associated with increased expression of *Gdf15* in the heart and increased levels of GDF15 in the circulation. We provide evidence that blockade of GDF15 activity prevents cachexia and slows the progression of heart failure. Our data suggests that cardiac stress mediates a GDF15 dependent pathway that drives weight loss and worsens cardiac function. We show relevance of GDF15 to lean mass and protein intake with patients with heart failure. Blockade of GDF15 could constitute a novel therapeutic option to limit cardiac cachexia and improve clinical outcomes in patients with severe systolic heart failure.

## Introduction

Heart failure (HF) is a common, complex condition with a poor prognosis and increasing incidence ^1^. Cachexia is associated with chronic heart failure and its occurrence independently predicts increased morbidity and mortality independently of other variables ^2^. The pathways that connect cachexia and heart failure are unclear. In particular, the extent to which impaired nutritional status might contribute to the deterioration of cardiac function needs investigation. The paucity of animal models of heart failure, that consistently develop cachexia, has impeded progress in this field.

The unfolded protein response (UPR) has been shown to have both protective and pathological roles in development of heart failure ^3 4^, and also feeding behaviour ^5 6^. The rapid adaptations to diverse stimuli that induce cytosolic and ER stress, such as protein misfolding, loss of metabolites, the production of reactive oxygen species and DNA damage, are in part instigated by the phosphorylation of (eIF2α). This evolutionarily conserved and vital cellular defence system enables tuning of the level of protein translation and a transcriptional reprogramming of the cell called the integrated stress response (ISR). Activation of the ISR leads to the induction of PPP1R15A (also known as GADD34), which binds protein phosphatase 1 (PP1) and G-actin to dephosphorylate P-eIF2α^7, 8^. PPP1R15A is positioned at a critical nexus in the ISR pathway where it participates in a negative-feedback loop, controlling levels of phosphorylated eIF2α and the activation of the ISR and protein translation.

Growth Differentiation Factor, GDF15, is an established biomarker of cellular stress and heart failure^9^. GDF15 acts through a receptor complex that is expressed solely in the hindbrain, through which it activates neuronal pathways that are perceived as aversive, and suppresses food intake ^10^. Reduced food intake has been shown to mediate most of the effects of GDF15 on body weight ^11 12 13^. Recently, GDF15 administration was shown to trigger conditioned taste avoidance in mice. GDF15 expression has been shown to be regulated by the ISR in response to nutritional stress ^14, 15^ but the contribution of PPP1R15A in this pathway has not been examined.

Although a clear biomarker of heart failure, it is not known whether GDF15 plays a contributory or protective role in this context. Here we examine the relationship of GDF15 with heart failure and its role in cardiac cachexia, utilising mice that lack catalytically active PPP1R15A^16^ In our initial studies examining the role of PPP1R15A in different models of heart failure using bone marrow transfer protocols involving whole body irradiation, we serendipitously identified that PPP1R15A is critical in the prevention of irradiation-induced left ventricular heart failure and associated cardiac cachexia. Hearts from mice lacking PPP1R15A show activation of the ISR, fibrosis, inflammation and high levels of *Gdf15* in heart and GDF15 in the circulation. We provide the first evidence that blockade of GDF15 activity not only prevents cardiac cachexia but remarkably also slows the worsening of cardiac function. Finally, we also provide evidence that suggests GDF15 may contribute to cachexia in patients with heart failure.

## Results

### Mice lacking functional PPP1R15A exhibit left ventricular dilated cardiomyopathy and severe weight loss following irradiation

Following standard protocols of whole body irradiation at 11 Gy in which mice are reconstituted with bone marrow (BM) immediately after irradiation to replenish the radiation sensitive immune cells, male mice lacking functional PPP1R15A, (*Ppp1r15a*^ΔC/ΔC^) exhibited an unusual response in that they exhibited a reduction in heart function at 6 weeks post irradiation. The reduction in heart function culminated in severe heart failure by 7-9 weeks after irradiation. Echocardiographic assessment of left ventricular fractional shortening (LVFS%) and heart geometry of the *Ppp1r15a*^ΔC/ΔC^ mice revealed the presence of a dilated cardiomyopathy, with a significant increase in LV internal diameters at systole and diastole and no increase in wall thickness (Figure S1a-e). The heart failure was also associated with cachexia, the severity of this model often requiring euthanasia due to loss of body weight or humane endpoints (Figure S1f).

These experiments were repeated with male *Ppp1r15a*^ΔC/ΔC^ and *Ppp1r15a*^+/+^ littermates. The response of the *Ppp1r15a*^ΔC/ΔC^ mice was highly reproducible as shown by the development of severe heart failure with an associated cachexia in all of the mice lacking functional PPP1R15A. Female *Ppp1r15a*^ΔC/ΔC^ mice showed an identical susceptibility with a slightly earlier time course. This response contrasted with WT littermates, which did not exhibit loss of heart function or decrease in body weight (Figure 1a-b). Comparison of % change in body composition from 4 weeks post irradiation to 9 weeks post irradiation (or earlier if mice reached human endpoints), using an independent cohort of *Ppp1r15a*^ΔC/ΔC^ and *Ppp1r15a*^+/+^ littermates showed both lean and fat mass were significantly decreased in *Ppp1r15a*^ΔC/ΔC^ mice (Figure 1c). Mice heterozygous for *Ppp1r15a* did not exhibit heart failure over the 22 weeks post-irradiation (Figure 1d). We also examined the role of functional PPP1R15A in BM-derived versus non BM-derived compartments. These data show that the BM genotype had no impact on the susceptibility of *Ppp1r15a*^ΔC/ΔC^ or the resistance of WT littermates to irradiation-induced heart failure or cachexia (Figure 1e,f) suggesting that loss of PPP1R15A activity in immune cells did not contribute to cardiac cachexia.

**Figure 1.**
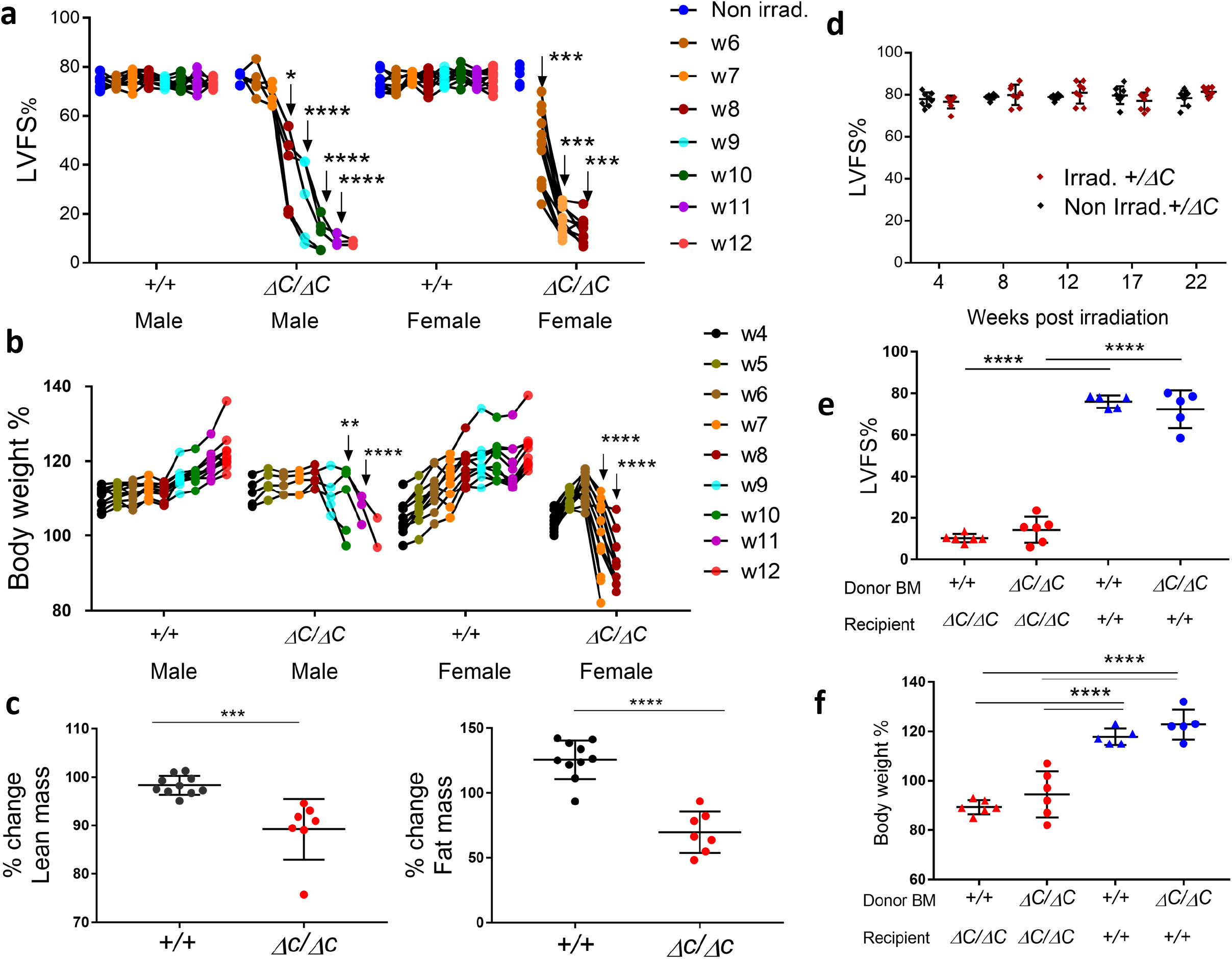
Mice lacking functional PPP1R15A exhibit severe heart failure and weight loss following whole body irradiation. Mice lacking functional PPP1R15A (^*ΔC/ΔC*^) n=5, or wild type female (^*+/+*^) n=10 and male (^*+/+*^) n=10 controls were subjected to whole body irradiation (11Gy) then bone marrow (BM) transfer and monitored by (**a**) echocardiography for left ventricular fractional shortening (LVFS%) and (**b**) body weight (% of body weight compared to day 0 of irradiation). The number of mice decreased over time due to euthanasia due to reaching humane endpoints. Male (^*ΔC/ΔC*^) and (^*+/+*^) mice are littermates. Female mice (^*ΔC/ΔC*^) n=10, shown from combined experiments, are not littermates. P values of data show comparison between genotypes at identical time points calculated by two-way ANOVA with Sidak’s correction for multiple comparisons. Male and female mice were calculated independently (**c**) TD-NMR analysis of body composition of lean and fat using an independent cohort of mice using two-tailed unpaired *t* test (**d**) Non-irradiated and irradiated *Ppp1r5a*^*+/ΔC*^ littermates heart function (LVFS%) up to 22 weeks. Irradiated *Ppp1r5a*^*+/+*^ *and Ppp1r5a*^*ΔC/ΔC*^ male mice reconstituted with either *Ppp1r5a*^*+/+*^ *or Ppp1r5* ^*ΔC/ΔC*^ BM, and show (**e**) LVFS% and (**f**) body weight at 9 weeks post-irradiation. e,f using two-way ANOVA with Tukey’s correction. **p < 0.01, ***p < 0.001, ****p < 0.0001.

### Heart failure in irradiated *Ppp1r15a*^ΔC/ΔC^ mice is associated with activation of the ISR

Given that the absence of functional PPP1R15A resulted in severe heart failure, we determined whether *Ppp1r15a* expression was induced in the hearts of irradiated WT mice. We analysed *Ppp1r15a* transcription using single molecule in situ hybridisation (SM-ISH), revealing basal expression of *Ppp1r15a* mRNA in non irradiated hearts that reached a maximal expression 5-7 weeks post-irradiation (Figure 2a,b and S2a). This peak was coincident with the detection of heart failure in mice lacking functional PPP1R15A.

**Figure 2.**
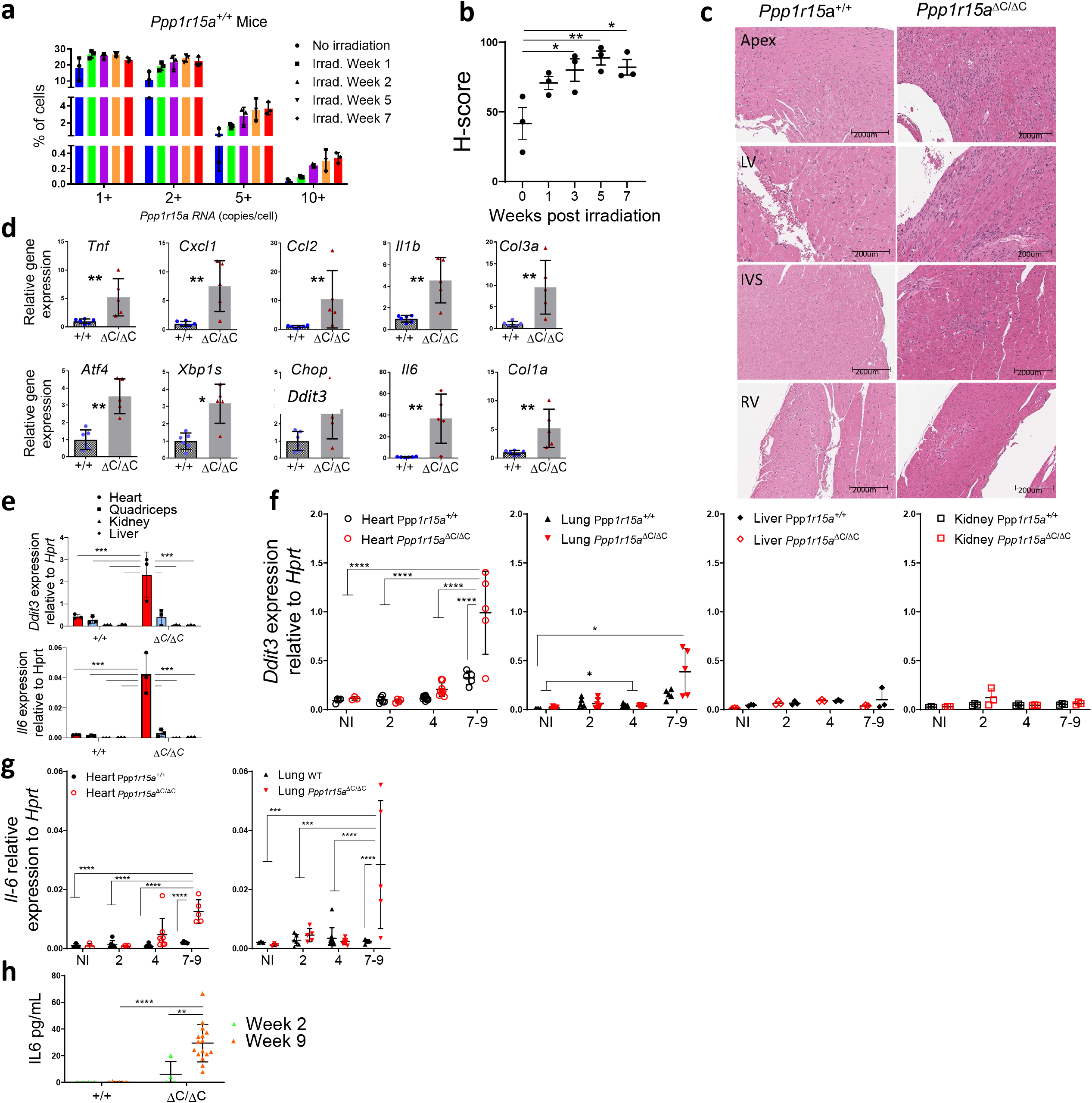
Heart failure is associated with local inflammation and activation of the integrated stress response. (**a**) *Ppp1r15a* transcript levels were assessed by Single Molecule In Situ Hybridisation (SM-ISH) in heart tissue sections derived from non-irradiated, or irradiated female WT mice, at 1,3,5 and 7 weeks post irradiation. (**b**) H-score of *Ppp1r15a* expression. H-score was analysed by ANOVA and Dunnett’s post test. (**c**) Analysis of tissue from *Ppp1r15a*^+/+^(^*+/+*^) or *Ppp1r15a*^*ΔC/ΔC*^(^*ΔC/ΔC*^) male mice (n=3 mice/group), 7 weeks post-irradiation with H&E staining (scale bar 200mm). (e) Heart, quadricep, kidney and liver tissue taken from male mice at 7 weeks from post-irradiation and analysed for expression for *Ddit*3 and *Il6* (n=3). Tissue taken from female mice at indicated time post-irradiation analysed for **(f)** *Ddit*3 (Heart, Lung, Liver. Kidney) and (**g**) *Il6* gene expression (heart and lung). 7-9 week samples taken at humane endpoint (h) IL6 in plasma taken at 2 or 9 weeks post irradiation or earlier if humane endpoints were exceeded. e,f,g were analysed by two-way ANOVA and Tukey’s post comparison All data expressed as mean +/-SD. *p < 0.05 **p < 0.01, ***p < 0.001, ****p < 0.0001.

In order to understand the reasons for the heart failure, we examined tissue sections taken from *Ppp1r15a*^ΔC/ΔC^ mice and WT littermates (^+/+^). Analysis of H&E stained sections of hearts from female mice at different time points post-irradiation, showed a progressive process of inflammation only in *Ppp1r15a*^ΔC/ΔC^ mice, evident from 5 weeks post-irradiation (Figure S2b). H&E stained sections of hearts from *Ppp1r15a*^ΔC/ΔC^ male mice, at 7 weeks post-irradiation, showed foci of myocardial inflammation, predominantly in the apex, LV and interventricular septum (IVS), with relatively few lesions within the right ventricle (Figure 2c). There were broad swathes of inflammation in which there was no recognisable myocardium, notably, neutrophils were inconspicuous on histology. These foci of inflammation were associated with fibrosis as confirmed by Sirius red staining and cross polarisation (Figure S2c, d). Immunostaining showed the inflammatory cells included CD3^+^ T cells and LAMP2^+^ macrophages (Figure S2e, f). These inflammatory changes were not evident in WT mice.

Analysis of transcript levels of the UPR, ISR, inflammatory and fibrotic markers in hearts of *Ppp1r15a*^ΔC/ΔC^ and WT littermates at 7 weeks post-irradiation revealed significant changes. Activation the ISR activation as indicated by upregulation of *Ddit3* and *Atf4*. Although *Atf4* gene expression can be mediated independently of eIF2α phosphorylation, *Ddit3* gene expression is dependent on eIF2αP and occurs as a result of activation of the ISR ^7^. The increase of XBP-1 splicing (*XBP1*s) indicated activation of the ER stress sentinel, IRE1, and the possible contribution of ER stress-mediated induction of UPR and ISR. Inflammatory genes *Tnf, Il1b, Ccl2, Cxcl1, Col1a* and *Col3a* were increased significantly, with the greatest fold change in *Il6*, confirming the histological findings of inflammation and fibrosis (Figure 2d).

Using *Il6* and *Ddit3* as indicators of inflammation and ISR activation, respectively, we analysed their expression in a panel of tissues, heart, quadricep, kidney and liver taken from *Ppp1r15a*^ΔC/ΔC^ and WT littermates at 7 weeks post irradiation (Figure 2e). ISR activation and *Il6* gene expression was upregulated in hearts of mice lacking PPP1R1A activity, and was not evident in skeletal muscle nor kidney or liver. In a separate cohort of female mice, heart, lung, kidney and liver tissue was harvested over the irradiation time course (Figures 2f,g). Activation of the ISR was evident at the late stage of the disease in the heart with a smaller induction in lung but not kidney and liver. *Il6* upregulation was also evident in heart and lung but not other organs tested. Given that IL-6 was also detected in plasma (Figure 2g), this suggests that local inflammation in the heart may also have systemic consequences.

### *Ppp1r15a*^ΔC/ΔC^ cardiomyocytes express high levels of GDF15 after irradiation

One candidate that may drive the cachexia in this model is a recently described peptide hormone, GDF15, whose induction by cellular stress and the ISR has been well described, and acts centrally in suppressing food intake^17, 18^. GDF15 can only mediate its effects centrally through increased levels in the circulation^19^. Therefore, we examined plasma GDF15 from *Ppp1r15a*^ΔC/ΔC^ and *Ppp1r15a*^+/+^ mice, non-irradiated, and at different time points post-irradiation. Plasma GDF15 was elevated following irradiation at 9 weeks post-irradiation in mice of both genotypes, although *Ppp1r15a*^ΔC/ΔC^ showed significantly higher levels (Figure 3a).

**Figure 3.**
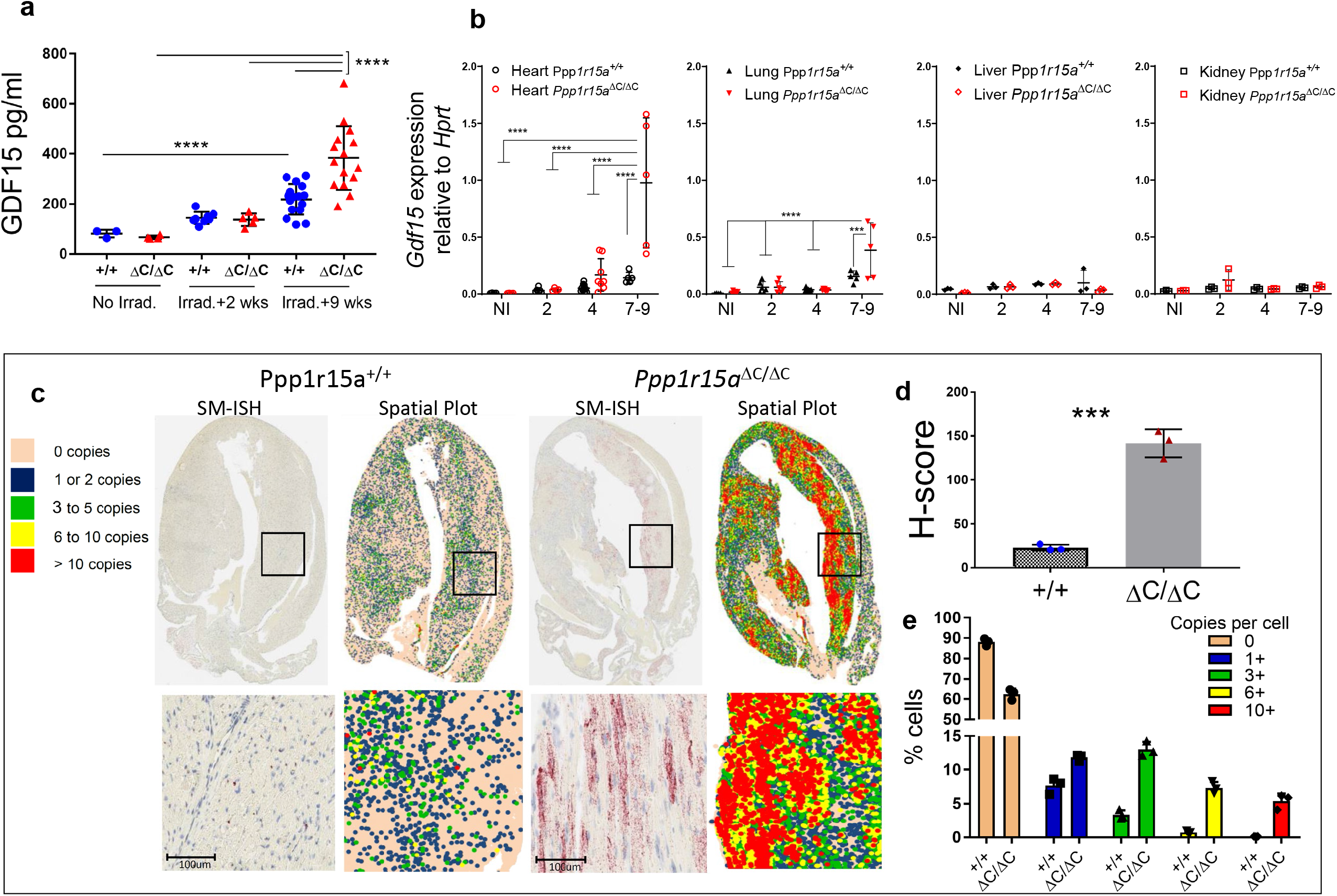
Absence of functional PPP1R15A results in increased expression of *Gdf15* following whole body irradiation. *Ppp1r15a*^*ΔC/ΔC*^(^*ΔC/ΔC*^) and *Ppp1r15a*^+/+^ (^*+/+*^) mice were irradiated (11Gy) and reconstituted with *Ppp1r15a*^+/+^ bone marrow. (a) GDF15 in plasma from non-irradiated mice, and at 2 and 9 weeks post-irradiation (or earlier if humane endpoints were exceeded. (b) *Gdf15* expression, by RT-qPCR, in heart, lung, liver and kidney taken at 7 weeks post-irradiation. *p* values by two-way ANOVA with Tukey’s correction. *Gdf15* transcript level in hearts, assessed by SM-ISH. (c) Representative spatial plots, (d) H-score and (e) and the distribution of cells expressing *Gdf15* transcripts, n=3 hearts/group. % cells and H-score are mean +/-SD (*P* values by students t test). ***p < 0.001, ****p < 0.0001.

In order to identify the source of GDF15 in the irradiated mice, we analysed *Gdf15* mRNA expression in heart, kidney and liver and lung, which identified the heart as potentially the major contributor to the increased GDF15 in plasma of *Ppp1r15a*^*ΔC/ΔC*^ mice (Figure 3b). To identify the cells expressing *Gdf15*, we utilised SM-ISH by RNAscope on tissue sections prepared from male *Ppp1r15a*^*ΔC/ΔC*^ and WT hearts at 7 weeks post-irradiation (Figure 3c). G*df15* mRNA was highly expressed in cardiomyocytes, identified by their morphology and presence of striations, in *Ppp1r15a*^*ΔC/ΔC*^ hearts, and significantly greater than irradiated WT mice when quantified by copies per cell and H-score (Figure 3d,e). *Gdf15*-expressing cardiomyocytes were abundant in apex, LV, and IVS, while relatively sparse in the right ventricles of *Ppp1r15a*^*ΔC/ΔC*^ mice. Typically, *Gdf15* expression was highest in peri-inflammatory foci of cardiomyocytes at 7 weeks, while inflammatory cells did not express conspicuous *Gdf15*. Analysis of hearts taken at different time points post-irradiation revealed a gradual increase in *Gdf15* expression over time, single cardiomyocytes expressing high quantities of *Gdf15* in the absence of inflammatory foci were detectable as early as 3 weeks post-irradiation (Figure S3). This suggests that cardiomyocyte GDF15 expression, an indicator of cellular stress, may precede the infiltration of the inflammatory cells in *Ppp1r15a*^*ΔC/ΔC*^ mice.

### The absence of CHOP does not prevent irradiation induced cardiac cachexia nor GDF15 expression

CHOP is a transcription factor highly induced during the UPR and deletion of the *Ddit3* gene has provided protection against tissue damage and disease, including atherosclerosis^20^ and renal damage caused by the UPR inducing toxin tunicamycin^21^. We therefore examined whether absence of *Ddit3* impacted on development of irradiation induced heart failure and weight loss. *Ppp1r15a*^*ΔC/ΔC*^ mice lacking *Ddit3* showed no protection from irradiation induced weight loss or heart failure (Figure 4a,b,c). CHOP has also been described as an important factor in inducing GDF15 expression following induction of the ISR ^22^ however, the absence of CHOP (*Ddit3)* had no significant impact in the levels of GDF15 detected in plasma nor expression in heart tissue (Figure 4e,f,g).

**Figure 4.**
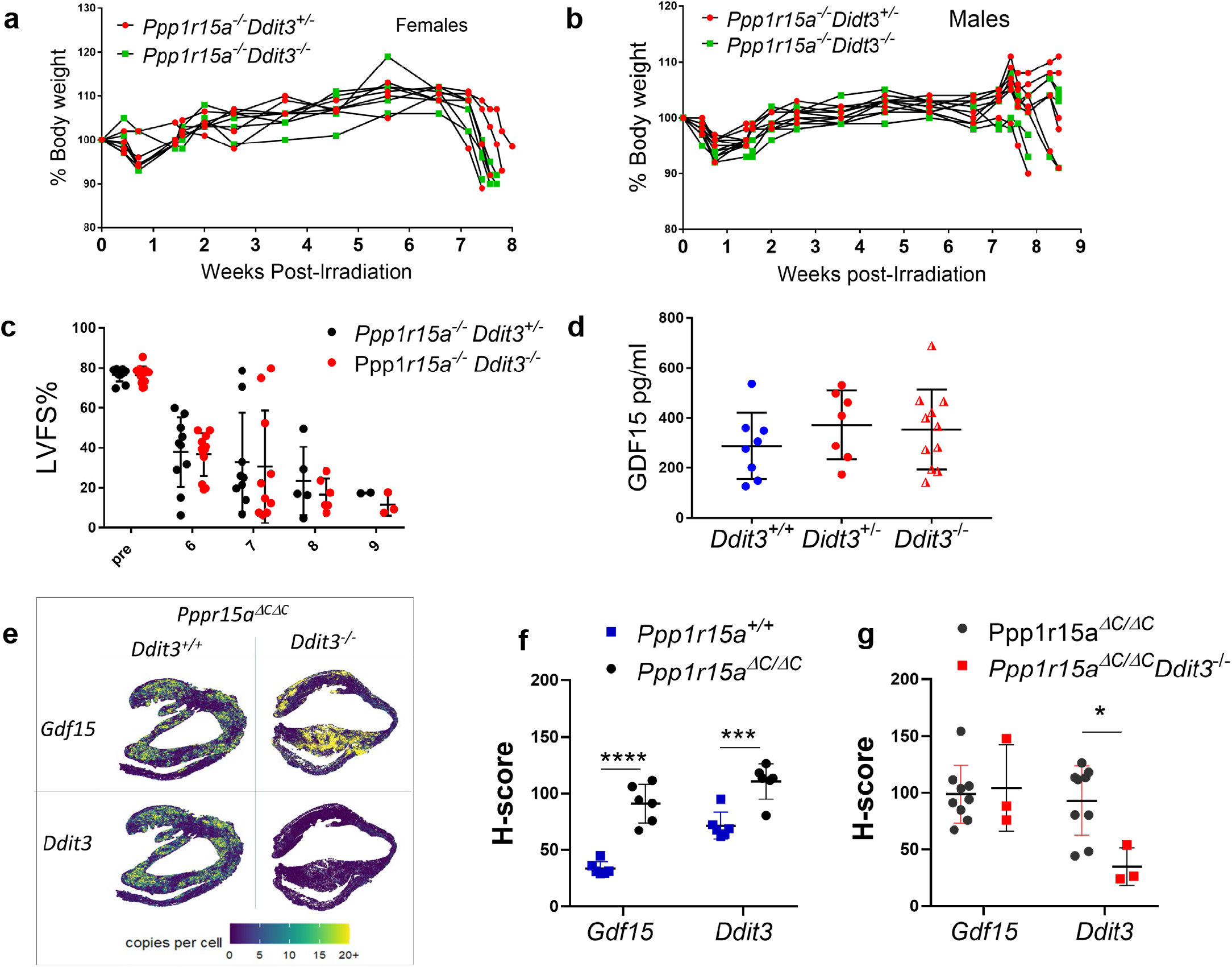
The absence of *Ddit3* does not prevent irradiation induced heart failure nor GDF15 expression. *Ppp1r15a*^*ΔC/ΔC*^ *Ddit3*^*+/-*^, *Ppp1r15a*^*ΔC/ΔC*^ *Ddit3*^*+/-*^ *or Ppp1r15a*^+/+^mice were irradiated (11Gy) and reconstituted with *Ppp1r15a*^*+/+*^ BM. Mice were culled at 8.5 weeks post irradiation or earlier due to humane end points. % of body weight of (**a**) female and **(b**) male mice. **(c)** LV function (LVFS %) assessed by echocardiography. **(d)** GDF15 analysis of the plasma. *Gdf15* and Ddit3 transcript levels in heart tissue assessed by SM-ISH and distribution shown in (**e**) spatial plots and (**f, g**) H-score of cells expressing *GDF15 and Ddit3* transcripts. H-score was analysed by ANOVA and Dunnett’s post test. *p* values determined by two-way ANOVA. Data show mean +/- SD. **p < 0.01, ***p < 0.001, ****p < 0.0001.

### Blockade of GDF15 activity prevents weight loss and cachexia and slows the worsening of cardiac function

Given the increased GDF15 in irradiated *Ppp1r15a*^*ΔC/ΔC*^ mice, we hypothesized that the cachexia was mediated by reduced food intake via GDF15 signalling. In order to test this hypothesis, we used a monoclonal antibody mAB2, against mouse GDF15. This monoclonal antibody has been validated to block the functional activity of GDF15, whereby mAB2 rapidly reversed the weight loss induced by recombinant GDF15 as opposed to mice receiving an isotype control^23^. To investigate whether GDF15 was driving cachexia we analysed whether mAB2 could inhibit loss of body weight in irradiated *Ppp1r15a*^**ΔC/ΔC**^ mice. mAB2 or IgG were administered at the beginning of week 4 post-irradiation and repeated at 3 day intervals until the end of week 9. Mice treated with control IgG exhibited severe weight loss from 8 weeks onward, which was in striking contrast to mice receiving mAB2 (Figure5a), suggesting that the weight loss in this model was GDF15-mediated. The loss in body weight in the control group was, in part, attributed to loss of fat mass, indicated by the lower gonadal fat pad weight in the IgG treated group (Figure 5b). The difference in severity of the treatment groups was highlighted by the finding that only mice in the IgG treated group required euthanasia, due to a combination of body weight loss and other welfare indicators (Figure 5c).

**Figure 5.**
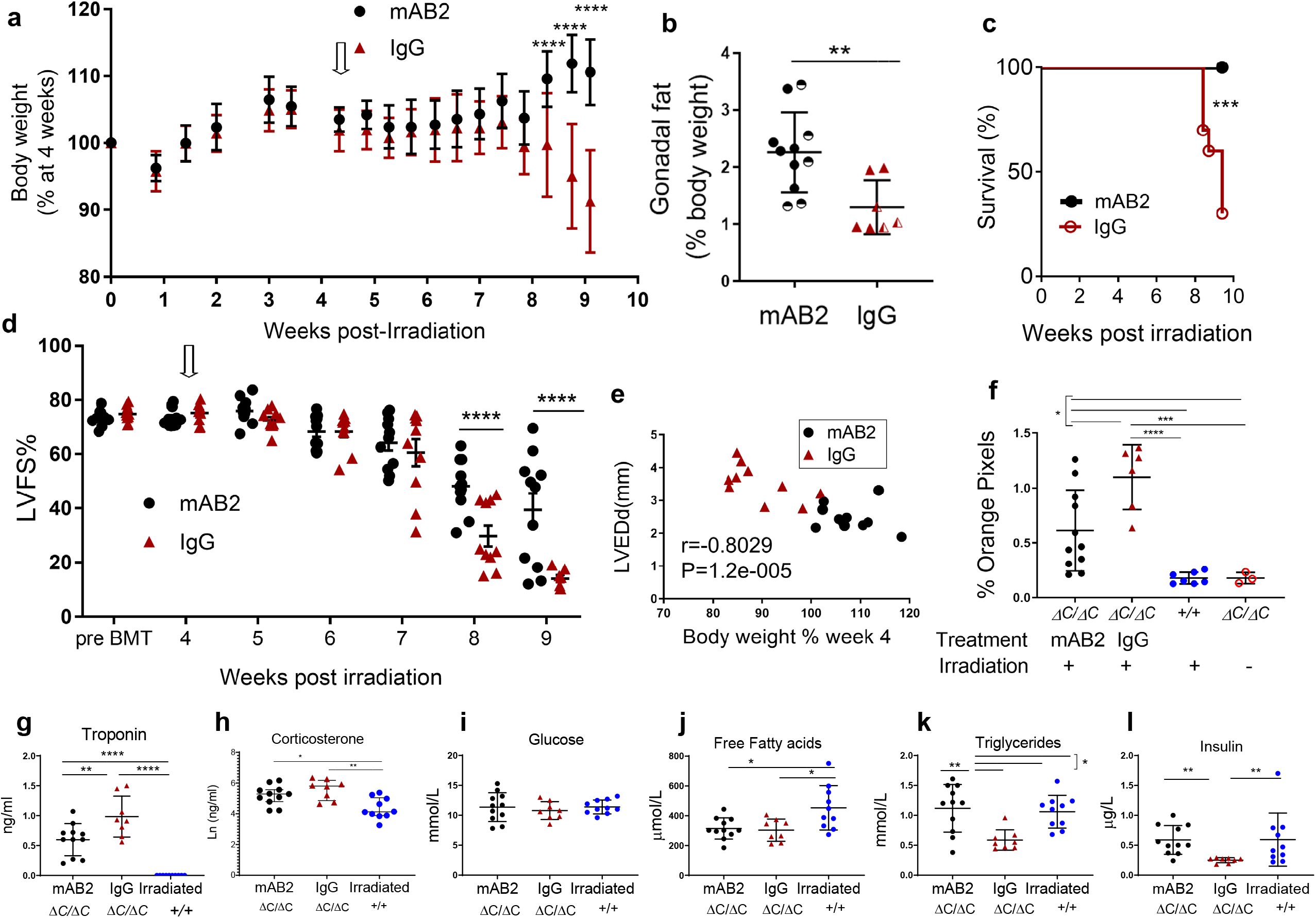
GDF15 is required for weight-loss and severe heart failure in irradiated mice lacking functional PPP1R15A. *Ppp1r15a*^*ΔC/ΔC*^ or *Ppp1r15a*^+/+^mice were irradiated (11Gy) and reconstituted with *Ppp1r15a*^*+/+*^ BM. At 4 weeks (open arrow), *Ppp1r15a*^*ΔC/ΔC*^ mice were randomly assigned and given an antibody that blocks GDF15 activity (αGDF15), n=11 or an isotype control antibody (Isotype Ab), n=10, at 10mg/kg, at 3-day intervals. Mice were culled at 9.5 weeks post-irradiation or earlier due to humane end points. (**a**) Body weights, % of body weight at 4 weeks post-irradiation. *p* values were determined by two-way ANOVA with Sidaks correction for multiple comparisons, p values indicated by asterisks, show comparison between treatment groups at identical time points. (**b**) Relative weight of fat pads, female mice indicated by half-filled symbols. p values were calculated by two-tailed Student’s *t*-test. (**c**) Survival plot, no mice were found dead but mice that approached humane endpoints were euthanised. Data analysed using Gehan-Breslow-Wilcoxon Test. (**d**) LV function (LVFS %) assessed by echocardiography. Data were analysed using ANOVA and Tukey’s post comparison. (**e**) Correlation of LV end diameter at diastole (LVEDd) with % body weight at 9 weeks post irradiation, r calculated by Pearson’s correlation coefficient. (**f**) Analysis of fibrosis in heart tissue sections using picrosirius red staining followed by microscopy under polarized light. Blood taken at 9.5 weeks post-irradiation (or earlier due to humane endpoints) and plasma analysed for (**g**) troponin and (**h-l**) metabolic parameters as indicated. Data (f,g,h,i,j,k) analysed using one way ANOVA and Tukey test post comparison. Data in (l) analysed using Kruskal-Wallis test with Dunn’s post test. All data show mean +/- SD. *p < 0.05 **p < 0.01, ***p < 0.001, ****p < 0.0001.

Unexpectedly, blocking GDF15 activity also attenuated the development of severe heart failure. GDF15 blockade significantly prevented the decrease in LV function compared to the IgG treated group (Figure 5d,e). This was also shown by differences in other parameters of heart function (Figure S4a-g). Heart fibrosis and plasma troponin a measure of cardiac damage, were also reduced by mAB2 (Figure 5f,g). The strong correlation of body weight with heart function suggests that these parameters are interconnected in this model (Figure 5e). Activation of the hypothalamic-pituitary-adrenal stress axis was evident by the increase in corticosterone in *Ppp1r15a*^ΔC/ΔC^ mice compared to irradiated WT mice (Figure 5h), which ruled out the possibility of adrenal gland dysfunction as a cause of weight loss. GDF15 has recently been shown to stimulate the hypothalamic-pituitary-adrenal axis in response to a range of stressful stimul^24^. Given that we show no significant difference in corticosterone levels in the presence of GDF15 blocking antibody in this mode, this would suggest that this property of GDF15 does not contribute to heart failure in this model. Analysis of other metabolic parameters (Figure 5i-l) showed that glucose levels in the blood were sustained, with insulin levels significantly reduced in IgG treated mice. Free fatty acids (FFAs) were significantly lower in *Ppp1r15a*^ΔC/ΔC^ compared to irradiated WT mice, with a reduction in plasma triglycerides only in IgG treated *Ppp1r15a*^ΔC/ΔC^ mice. These data suggest that GDF15 signalling pathways contribute towards cardiac cachexia in this model and impact significantly on the development of the dilated cardiomyopathy.

### GDF15 expression correlates with parameters of heart function in another murine model of cardiac cachexia

We also examined GDF15 expression in a murine model of dilated cardiomyopathy where cardiac-specific ablation of Yme1l (cYKO) in mice induces mitochondrial fragmentation and altered cardiac metabolism. These mice have been previously shown to exhibit dramatic weight loss at approximately 42 weeks of age^25^. We measured plasma GDF15 in a cohort of these mice that exhibited a reduction in heart function at 34-35 weeks of age, at this stage, it was not expected that all mice would be at the final stage of the disease, although a proportion of the mice exhibited rapid weight loss (Figure S5a,b). Plasma GDF15 correlated strongly with changes in parameters of heart geometry and heart function (Figure S5c-e).

### GDF15 expression correlates with cachexia in a cohort of patients with heart failure

Weight loss is strongly associated with adverse outcomes in patients with chronic heart failure ^26^. In addition, GDF15 has been identified as an important predictor of mortality in patients with heart failure^9^. To investigate the association between GDF15 and cardiac cachexia we analysed The BIOlogy Study to TAilored Treatment in Chronic Heart Failure (BIOSTAT-CHF) cohort, which included 2516 patients with worsening signs and/or symptoms of heart failure ^27^. Protein intake in 24-hour urine was calculated by the Maroni formula. Since spot samples were used, the adjusted Maroni formula was utilised. Baseline levels of GDF-15 were inversely correlated with protein intake (Figure 6a). The concentration of serum creatinine, produced after stable conversion of creatine largely found in skeletal muscles, can also be used as a marker to reflect (peripheral) muscle catabolism and is an established marker of muscle mass ^28^, and can be used as a measure of cachexia^29, 30^. The adjusted formula and use of spot urine samples was validated in a heart failure population using data from the Additive renin Inhibition with Aliskiren on renal blood flow and Neurohormonal Activation in patients with Chronic Heart Failure and Renal Dysfunction cohort (ARIANA-CHF-RD)^31^. GDF15 inversely correlated with urinary creatinine (Figure 6b). These associations remained statistically significant for both protein intake (standardized β: −0.20, P <0.001) and urinary creatinine (β: −0.18, P <0.001) in multivariable regression analyses mutually adjusted for age, sex, BMI at baseline, log transformed eGFR and a medical history of diabetes. Furthermore, patients that fitted the criteria for cachexia, exhibited greater plasma GDF15 than those that did not fit the criteria (Figure 6c).

**Figure 6.**
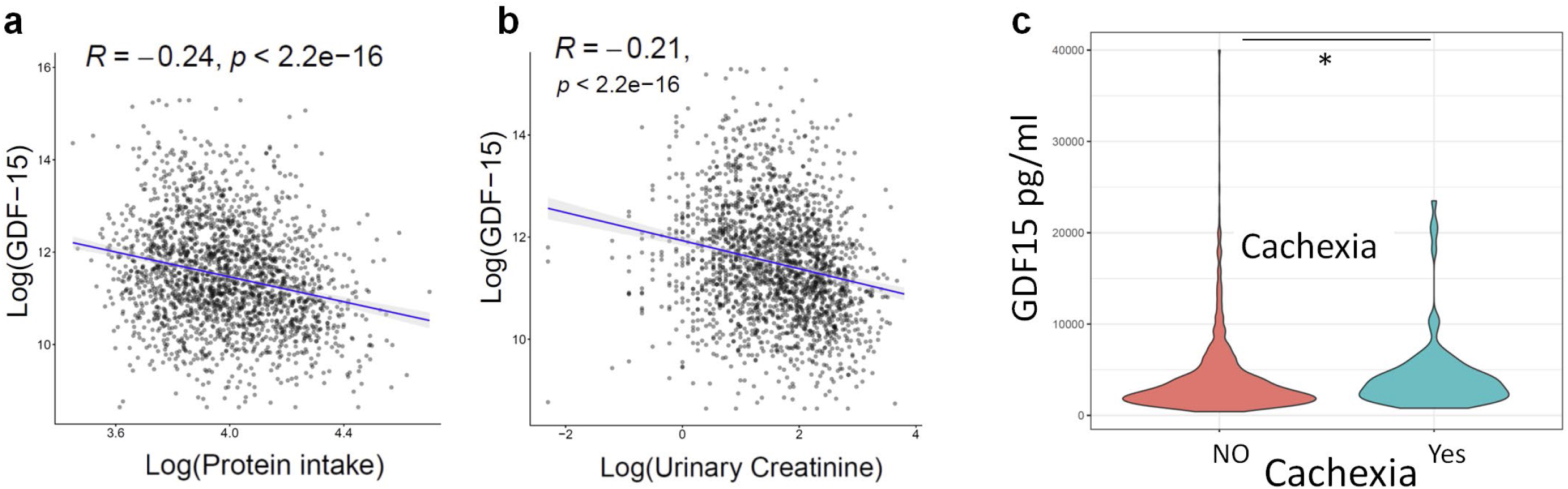
GDF15 correlates with parameters of cachexia in patients with heart failure. Analysis of the BIOSTAT-CHF cohort of patients with heart failure, correlation between Log transformed plasma GDF-15 and (**a**) Log transformed estimated protein intake and **(b)**, Log transformed urinary creatinine. (**c**) Violin plots of plasma GDF15 compared in patient groups according to whether they fit the criteria for cachexia. Data analysed using Mann-Witney U test *p < 0.05.

## Discussion

We identified PPP1R15A as an essential factor in protection from radiation-induced heart failure and associated cardiac cachexia. The mice lacking functional PPP1R15A exhibited a dilated cardiomyopathy histologically characterised by loss of cardiomyocytes, a prominent immune cell infiltration, fibrosis and pro-inflammatory cytokine expression.

The presence of inflammatory infiltrates at 4 weeks post irradiation suggest that this is a precursor to heart dysfunction in mice lacking PPP1R15A activity. Cardiomyopathy developed irrespective of the PPP1R15A status of BM-derived cells, indicating that the susceptibility to cardiomyopathy was intrinsic to the heart. The finding that the inflammation, *Gdf15* and ISR activation predominated in heart and not skeletal muscle nor kidney and liver suggests that heart tissue is intrinsically vulnerable to irradiation in the absence of PPP1R15A. GDF15 expression is associated with mitochondrial dysfunction and cellular stress and is clearly evident in the cardiomyocytes of *Ppp1r15a*^*ΔC/ΔC*^ mice at 7-9 weeks post irradiation. The cardiomyocyte cellular stress may be the primary dysfunction or secondary to perturbation of other cell types, such as capillary endothelial cells which are radiation sensitive^32^.

PPP1R15a functions in its canonical role to limit the extent of ISR activation and reversal of stress-induced translational attenuation^7, 33, 34^. In support of this, we show evidence of enhanced ISR activation in *Ppp1r15a*^*ΔC/ΔC*^ mice in heart tissues but this only occurred at 7-9 weeks post irradiation when weight loss and heart dysfunction was evident. This suggests that a pivot point may exist where the loss of PPP1R15A and dysregulated ISR drives cell death and an inflammatory cascade. In tissue culture PPP1R15A (GADD34) mutant cells are more prone to cell death in response to ER stress induced by thapsigargin^7^. This agent causes ER stress and induction of the UPR and ISR through depletion of ER calcium stores and is particularly effective at shutting down protein synthesis, indicating that recovery of protein translation is necessary for the adaptive pro-survival effects.

This protective role of PPP1R15a shown here is counter to studies where mice lacking PPP1R15a are protected from UPR-induced nephrotoxicity^16^ suggesting that specific organs may have different requirements for sustaining protein synthesis in the face of ISR activation. The use of small molecules that mimic the effect of PPP1R15A activity by preventing translational shut down^35^, are attractive candidates to utilise *in vivo* to test the hypothesis that PPP1R15A activity protects WT mice from heart failure by preventing over-exuberant translation attenuation and ISR activation.

One consistent feature in the irradiated *Ppp1r15a*^*ΔC/ΔC*^ mice was the prominent fibrosis of the left ventricle at the later stages of the disease. A previous study examining wound healing, showed that mice lacking PPP1R15a exhibited accelerated wound closure, increased number of myofibroblasts, and elevated collagen production^36^. A recent report has also shown that mice lacking *Ppp1r15a* show increased pulmonary fibrosis in response to the DNA damage inducing chemical bleomycin^37^. Taken together, these data suggest that the PP1R15A may play an important role in the negative regulation of fibrosis pathways.

Although CHOP has been shown to sensitize cells to the toxicity of ER stress and UPR induction, its absence protects in a number of different models of disease ^20, 38, 16^, we show no role for CHOP in this model of cardiac cachexia. It was surprising that the absence of CHOP (*Ddit3*) in *Ppp1r15a*^*ΔC/ΔC*^ mice did not impact the expression of GDF15 in cardiac tissue. CHOP has been shown to act cooperatively with ATF4 to regulate Gdf15 expression induced by ISR activation^14, 22, 39^. It is plausible that other factors in combination with ATF4, contributes to GDF15 expression in the absence of *Ddit3* in heart tissue of irradiated *Ppp1r15a*^*ΔC/ΔC*^ mice. p53 has been shown to be an important driver of GDF15 expression and cachexia in toxin and DNA damage models and may be a critical factor in this model of cardiac cachexia ^40 41, 42^. Given that non irradiated *Ppp1r15a*^*ΔC/ΔC*^ mice do not show any cardiac dysfunction, we propose that DNA damage in combination with a dysregulated ISR are critical factors in the development of GDF15 medicated cardiac cachexia. Although This raises the question whether susceptibility to heart failure following cancer therapies involving DNA damaging reagents such as anthracyclines and cisplatin result from a dysregulated ISR and GDF15 expression.

The protective effects of anti-GDF15 treatment in heart failure are in contrast to the finding that GDF15-deficient mice were more susceptible to cardiac rupture in a murine model of acute myocardial infarction^43^. However, it was unclear whether the cardiac rupture was precipitated by the absence of GDF15 given that no data was provided to show that cardiac rupture could be prevented by GDF15 supplementation. More recently, Luan et al found that anti-GDF15 treatment resulted in increased cardiac damage following injection of LPS over 48 hours^44^. A caveat of that study was that an antihuman GDF15 was used to block endogenous murine GDF15, while some evidence was presented showing its ability to attenuate murine GDF15 signalling, no data was provided to show the efficacy of this antibody in the prevention of weight loss induced by the rGDF15. These findings are contradicted by a study using a validated blocking antibody raised against murine GDF15, where no effect was shown in lethality or weight loss following high or low dose LPS treatment^23^.

The hindbrain receptor for GDF15 has recently been identified and its action by ligand powerfully suppresses food intake^11 12 13^. The neutralisation of GDF15 in our model substantially slowed the development of weight loss. We assume this occurred by a mechanism that at least in part involved an improvement in food intake, although we were unable to undertake single housing experiments necessary to measure this directly. Our analysis of metabolites in the circulation suggests that the most cachectic mice had significantly lower nutrient stores and that blocking GDF15 maintains food intake and triglyceride levels. Strikingly, inhibition of GDF15 activity had a substantial impact on the development of severe heart failure and the necessity for euthanasia. We show that blocking GDF15 reduced damage to the heart but also reduced heart fibrosis, and we presume this protection occurs down stream of GDF15 signalling in the hind brain although the pathway for this protection needs further investigation. It is not known whether the loss of GDF15 signalling mediates its cardio-protection via its effects on body weight, intake of food that provides vital fuel substrates for the heart, or additional signalling circuits emanating from the hindbrain that mediate cardio-protective effects. Irradiated wildtype mice also showed a significant increase in plasma GDF15 albeit at a lower amount than mice lacking Ppp15r15a activity. This may suggest that that both the context and concentration of GDF15 are important factors driving cardiac cachexia. Nutritional stress resulting from reduced food intake may compound and exacerbate cardiomyocyte dysfunction resulting from irradiation in the absence of PPP1R15A activity. The strong correlations of GDF15 in plasma and parameters of heart function in the mice lacking *Yme1l* and exhibiting dilated cardiomyopathy, suggests that GDF15 may have broad applicability in driving the severity of heart failure in cardiac cachexia.

In a recent rat model of monocrotaline induced right-ventricular dysfunction, blocking GDF15 restored the ability of rats to gain weight over the treatment period but had no impact on heart function. Since the rats did not lose weight over the experimental period, this model was not able to test the effect of blocking GDF15 on heart function in the context of cardiac cachexia^45^.

Using a cohort of patients with heart failure, we showed that GDF15 measured in plasma correlated with surrogate markers in urine for protein intake and muscle mass. Although the study utilised spot urine samples rather than a 24 hr urine sample, our recent study using spot urine analysis showed that urinary creatinine concentrations were associated with smaller body dimensions (lower BMI, height, and weight) and an increased risk of (*>*5%) weight loss at 9 months^31^. The inverse correlation of muscle mass and protein intake with GDF15 and raised GDF15 levels in patients with cachexia, support the concept that GDF15 plays a role in driving cardiac cachexia. Further analyses would be required to determine if GDF15 reduced protein appetite specifically or whether this reduction reflected a reduction in total food intake. In addition more in-depth analyses may also reveal the relationships between changes in GDF15 levels and changes in parameters of cachexia and heart remodelling.

In summary, our work identifies GDF15 as a critical driver of cardiac cachexia and critically impacted on heart function. This warrants further studies to investigate its role in human heart failure. Blockade of GDF15 could constitute a novel therapeutic option to limit cardiac cachexia and improve cardiac function and clinical outcomes in patients with severe systolic heart failure.

## Methods Animal studies

Mice were maintained in a 12 h:12 h light:dark cycle (lights on 07:00–19:00), temperature-controlled (22 °C) facility, in specific pathogen free conditions, with ad libitum access to food (RM3(E) Expanded Chow (Special Diets Services) and water. Sample sizes were determined on the basis of homogeneity and consistency of characteristics that were sufficient to detect statistically significant differences in body weight, food intake and serum parameters between groups. Experiments were performed with animals of both genders.

The *Ppp1r15a*^*ΔC/ΔC*^ mouse (originally named *Ppp1r15atm1Dron*)^16^ had been previously backcrossed to achieve 97% C57BL/6 purity as described ^5^. Experimental cohorts of male *Pppr15a* ^*ΔC/ΔC*^ and wild-type mice were generated by het × het breeding pairs unless otherwise stated. Data in Figure S1 are from *Ppp1r15a*^*ΔC/ΔC*^ mice irradiated at 11-12 weeks old. Data in Figure 1 are from *Ppp1r15a*^*ΔC/ΔC*^ male mice and male and female wildtype littermates irradiated at 7-8 weeks old. Data in Figure S2A are from female *Ppp1r15a*^*ΔC/ΔC*^ and wildtype littermates irradiated at 10-11 weeks old. Data shown in figure 2AC is from wildtype C57BL/6 mice irradiated at 9 weeks old.

### Irradiation and bone marrow transfer

Mice were maintained overnight with Baytril before irradiation were subject to whole body irradiation with two doses of 5.5 Gy (separated by 4 h) followed by reconstitution with 1× 10^7^ bone marrow (BM) cells. Baytril was administered for 4 weeks after bone marrow transfer. Mice were monitored by body weight and echocardiography over the experimental period. *Ppp1r5a* ^*ΔC/ΔC*^ and wildtype mice were euthanized by CO_2_ asphyxiation at the desired time point or for a maximum of 12 weeks. Mice were culled earlier if humane endpoints were approached. Blood was obtained from Vena Cava immediately at sacrifice, plasma separated by centrifugation at 6000xg and stored immediately at −80°C. Isolated organs were fixed in 4% paraformaldehyde for 24 hours at room temperature and then stored in 70% ETOH at 4°C prior to being embedded in paraffin. Alternatively, tissues were fresh frozen on dry ice and kept at −80 °C until the day of RNA extraction.

For the study involving the use of anti-GDF15 and isotype control antibodies, 13-14 week old *Ppp1r5a*^+/+^ *or Ppp1r5a* ^*ΔC/ΔC*^ male and female littermates were irradiated with 11Gy and reconstituted with sex matched *Ppp1r5a*^+/+^ BM. At 4 weeks post irradiation, *Ppp1r5a* ^*ΔC/ΔC*^ male and female mice were randomized into the treatment groups with n=5 of each sex in Isotype Ab group and n=5 females, n=6 males in the αGDf15 treated group. The mice were assigned by an independent researcher on the basis of body weight such that the mean body weights of each group were not significantly different. Irradiated *Ppp1r5a*^+/+^ mice were not given any treatment and were monitored by echocardiography only at 9 weeks post-irradiation prior to cull and harvesting of tissue. mAB2 and IgG isotype control were prepared as described for the cancer cachexia model and administered via S.C. injection once every 3 days from 4 weeks post-irradiation, which was repeated every 3 days until the end of the experiment. Researchers who administered the anti-GDF15 and isotype control antibodies were blinded to the identity of the antibody treatment which was prepared by an independent researcher Mice were monitored by echocardiography prior to irradiation and weekly from 4 weeks post-irradiation. The capture and analysis of echocardiography data was performed by a researcher that was double blinded to both group allocation and identity of the treatments. *Ppp1r5a* ^*ΔC/ΔC*^ mice were euthanized at 9.5 weeks post-irradiation or earlier if mice approached humane endpoints, determined by body weight loss and general welfare indicators.

These endpoints were assessed by a researcher who was blinded to the assignment of the treatment groups. After euthanasia, blood was immediately obtained via the Vena Cava and tissues harvested. Gonadal fat pads were dissected and weighed, organs were fixed immediately in 4% paraformaldehyde for 24 hours at room temperature and then stored in 70% ETOH at 4°C.

#### Echocardiography

Two-dimensional and M-mode echo were employed to detect the wall motion, the chamber dimensions, and the cardiac function. Cardiac function was assessed on conscious, un-anaesthetised mice with the Vevo 3100 ultrasound system (VisualSonics, 30 MHz MX400 probe). Echocardiographic M-mode images were obtained from a parasternal short axis view and cardiac function was measured on M-mode images. From two-dimensionally targeted M-mode tracings, left ventricular (LV) end-diastolic (LVEDd) and end-systolic diameter (LVESd), and LV anterior wall thickness (LVAWd) and posterior wall (LVPWd) dimensions were measured using Vevolabs software (Visual Sonics). Cardiac contractile function was assessed by LV fractional shortening (LVFS%), which was calculated as follows: LVFS(%) = (LVEDd – LVESd)/ LVEDd)× 100. LV mass was calculated LV Mass (mg) =1.053 x [(LVIDd+LVPWd+IVSd)^3^-LVIDd3]

#### Excluded samples

No sample or animal data were excluded from the data sets unless there was a collection failure. 3 fat pad data points are absent from isotype Ab data set (Figure 4B) as these mice had deteriorated rapidly and were culled by a technician who did not remove and weigh the fat pads. Data from 2 plasma samples were excluded from isotype Ab treated data set (Figure S4 H-L), due to inconsistent sample collection where 2 blood samples were left overnight at room temperature rather than an immediate plasma isolation and −80°C freeze.

### Dilated cardiomyopathy model using *Yme1l*^*-/-*^ mice

The cardiac specific *Yme1l*^*-/-*^ mice (cYKO) were generated by cross breeding *Yme1l*^*LoxP/LoxP*^ flanking exon 3 mice with mice expressing Cre-recombinase under control of the *Myh6* promoter as previously described ^25^. Adult mice were maintained under pathogen-free conditions in a temperature-controlled room and a 12-hour light-dark cycle at the CNIC animal facility. Chow diet and water were available ad libitum. Animal experiments conformed to European Union Directive 2010/63EU and Recommendation 2007/526/EC, enforced in Spanish law under Real Decreto 1386/2018. All experiments were approved for CNIC ethics committee and the Animal Protection Area of the Comunidad de Madrid (PROEX 176.3/20). Echocardiography was performed as above but using Vevo 2100.

### SM-ISH RNAscope

Detection of mouse *Gdf15* and *Ppp1r15a* transcripts was performed on Formalin-Fixed Paraffin-Embedded (FFPE) sections using Advanced Cell Diagnostics (ACD) RNAscope® 2.5 LS Reagent Kit-RED (Cat No. 322150), RNAscope® LS 2.5 Probe-Mm-Gdf15 (Cat No. 318528) or RNAscope® LS 2.5 Probe-Ppp1r15a (Cat No. 556038) (ACD, Hayward, CA, USA). Briefly, sections were cut at 10μM thick, baked for 1 hour at 60°C before loading onto a Bond RX instrument (Leica Biosystems). Slides were deparaffinized and rehydrated on board before pre-treatments using Epitope Retrieval Solution 2 (Cat No. AR9640, Leica Biosystems) at 95°C for 15 minutes, and ACD Enzyme from the LS Reagent kit at 40°C for 15 minutes. Probe hybridisation and signal amplification was performed according to manufacturer’s instructions. Fast red detection of mouse *Gdf15* and *Ppp1r15a* was performed on the Bond Rx using the Bond Polymer Refine Red Detection Kit (Leica Biosystems, Cat No. DS9390) according to the ACD protocol. Slides were then removed from the Bond Rx and were heated at 60°C for 1 hour, dipped in Xylene and mounted using EcoMount Mounting Medium (Biocare Medical, CA, USA. Cat No. EM897L). The slides were imaged on the Aperio AT2 (Leica Biosystems) to create whole slide images. Images were captured at 40x magnification, with a resolution of 0.25 microns per pixel.

### Spatial plots

*Ppp1r15a;* Cells with an expression score of 1 or more cells were selected and used to generate spatial plots in HALO, in which individual spots represent single cells coloured by expression level from blue (low expression) to red (high expression). *Gdf15*; Cells expression scores were separated into 5 bins of 0, 1-2, 3-5, 6-10 and greater than 10 spots per cell. These data were displayed in HALO using a colour key for the different bin sizes.

### Tissue processing and RNA analysis

The fresh heart, liver and kidney were harvested and snap frozen in liquid nitrogen. Tissue samples frozen at −80°C were handled on dry ice and 20mg of the tissue was placed in a pre-cooled 1.5mL flat-bottom tube containing 4 Yttria-stabilised (Y57) Grinding Media 5mm Balls (Inframat Advanced Materials) and 600uL of Buffer RLT from Qiagen RNeasy Mini Kit. Tissue was subsequently homogenised in a Qiagen^®^ Tissuelyser LT for 4 minutes at 50Hz and lysate transferred to 1.5Ml rounded-bottom Eppendorf tube. Qiagen RNeasy Mini Kit (no. 74106). Purified RNA solutions were then quantified by absorbance at 260nm using a Nano drop device.

For Figure 2C, 500ng of RNA was reverse-transcribed using QuantiTect Rev. Transcription Kit (Qiagen) according to manufacturer’s instruction. Real-time PCR was performed by using SYBR Green qPCR mix (Eurogentec) on a Roche Lightcycler.

For Figure 2D and 3C, 500ng of RNA was used to generate cDNA using Invitrogen SuperScript VILO cDNA Synthesis Kit (Catalog number:11754050) according to manufacturers instructions. Gene expression was analysed via TaqMan™ RNA-to-CT™ 1-Step Kit (catalog number: 392653) 2× universal PCR Master mix (Applied Biosystems Thermo Fisher, 4318157) and analysed using 7500 Real time Fast, PCR machine (Applied Biosystems,Thermo Fisher).

All reactions were carried out in either duplicate or triplicate and *C*_t_ values were obtained. Relative differences in gene expression were normalized to the expression levels of the housekeeping genes *Hprt or 36B4*. All qPCR primers are shown in the key resources table.

### Immunofluorescence

Paraffin (PFA) sections (stored at 4°C) were dried for 30 minutes, before being rehydrated in PBS for 10 minutes before the staining. PFA-fixed sections were then permeabilised in 0.1 % Triton X-100, 0.1 % Citrate buffer pH 6.0 (Dako) for 30 minutes. They were then washed in PBS, and incubated with the blocking solution (flow buffer + 5% serum of secondary antibody species, i.e. goat or donkey) for 30 minutes before being incubated with primary antibodies diluted in the blocking solution at indicated concentrations (See extended data table) overnight at 4°C. Samples were extensively washed with PBS and incubated with secondary antibodies diluted in the blocking solution at indicated concentrations (shown in key resources table) for 4 hours. Samples were again washed extensively in PBS, nuclei were counterstained with Hoechst 33342 (Invitrogen) and samples were mounted with CC mountTM (Sigma). Interstitial collagen was detected using picrosirius red staining followed by microscopy under polarized light.

### Measurement of parameters in plasma

In heart failure studies, GDF15 levels were measured using Mouse/Rat GDF15 Quantikine ELISA Kit (no. MGD-150, R&D Systems). In cancer studies GDF15 levels were determined using the mouse specific ELISA (R&D Systems, Minneapolis, MN). Serum IL6 levels were quantified using an Invitrogen eBioscience Mouse IL-6 ELISA Ready-SET-Go Kit (15511037) according to manufacturers instructions. Plasma Troponin-I level was measured using the Muscle Injury Panel 3 Mouse Kit (cat# K15186C, Meso Scale Diagnostics). Plasma non-esterified fatty acid concentration was measured using the Roche Free Fatty Acid Kit (half-micro test) (cat#11383175001, Sigma Aldrich). Plasma levels of glucose and triglycerides were measured on a Dimension EXL Analyser (Siemens Healthcare, Erlangen, Germany) using the DF30 and DF69A cartridges (Siemens Healthcare), respectively. Plasma concentration of insulin was measured using the Mouse/Rat Insulin Kit (cat# K152BZC-3; Meso Scale Diagnostics). Plasma corticosterone level was measured by the IDS Corticosterone EIA kit (Immunodiagnostic Systems).

## HUMAN STUDIES

### Patient population

BIOSTAT-CHF was an international, multinational, observational study, of which design and primary results have been published previously^27^. Briefly, patients were included in the in- or outpatient setting, received ≤50% of target dosages of ACEi/ARB and/or beta-blockers at time of inclusion and were anticipated to be uptitrated by the treating physicians. Patients were required to have a left ventricular ejection fraction (LVEF) of ≤40% at inclusion or have plasma concentrations of BNP and/or NT-proBNP >400 pg/mL or >2000 pg/mL respectively. Out of 2516 patients from the original BIOSTAT-CHF study, GDF-15 levels were available from 2300 patients.

### Measurements

Plasma concentrations of GDF-15 were measured using electrochemiluminescence on a cobas e411 analyzer, using standard methods (Roche Diagnostics GmbH, Mannheim, Germany) at baseline. Data on demographics including age, sex, medical history and weight and height were captured for all patients at baseline. Renal function was assessed using the estimated glomerular filtration rate (eGFR) based on serum creatinine measured at baseline. Urine samples were available in 2282 patients from the index cohort. Baseline spot sample urine measurements were stored at −80°C. Protein intake in 24-hour urine was calculated by the Maroni method, which was adjusted for the use of spot samples as previously published ^46, 47^. The formula used for protein intake in gram/day was as follows: 13.9 + 0.907 * Body mass index (BMI) (kg/m2) + 0.0305 * urinary urea nitrogen level (mg/dL). Urinary creatinine was determined at baseline in spot-urine samples of 2144 patients. Patients fit the criteria for cachexia if BMI was less than 20 kg/m2 and exhibited at least one of the following, CRP>5mg/ml, Hb<12g/dL or Albumin <3.2g/dL^48^.

### Statistical analyses

To investigate the association between baseline levels of GDF-15 urinary creatinine and protein intake as a measure of cachexia, we performed univariable continuous regression with log transformed GDF15 as the independent variable and urinary creatinine, estimated protein intake, as the independent variables. In subsequent analyses, these were mutually adjusted for possible confounders including age, sex, BMI at baseline, log transformed eGFR and a medical history of diabetes. All analysis is performed using R (version 3.5).

### Statistical Analysis

All numeric data were analysed using Graphpad Prism (USA). H-score was calculated = Σ (bin number x percentage of cells per bin). Data are represented as arithmetic mean ± SEM or show individual data points with arithmetic mean +/-SD. Differences between values were examined using the parametric two-tailed unpaired Student’s t test. Means of multiple groups were compared by one way or two way ANOVA and corrections made for multiple comparison using either Tukey’s, Sidak’s or Dunnet’s post tests which was dependent on the comparisons being made and were assigned by the PRISM analysis software. All tests were two-tailed. P values <0.05 were deemed significant.

### Study Approval

All studies in U.K. were performed in accordance with UK Home Office Legislation regulated under the Animals (Scientific Procedures) Act 1986 Amendment, Regulations 2012, following ethical review by the University of Cambridge Animal Welfare and Ethical Review Body (AWERB)

## Author contributions

Overall conceptualization by J.C.G. and Z.M. S.O.R, M.T, D.B. A.A.V also contributed to conceptualization of individual experiments and studies included in this body of work. Experimental investigation and data analysis by J.C.G, M.T, D.R, S.J, J.A.T, J.T, A.H, S.A, M.A, J.H, R.V.G, E.O.P, B.I and A.P.C. The paper was written by J.C.G, which was reviewed and edited by M.T, S.J.M, A.A.V, S.O.R and Z.M.

## Acknowledgments

Z.M. is supported by the British Heart Foundation professorship and grants (35TCH/10/001/2764235T, 35TRG/15/11/3159335T, and 35TPG/17/9/3283435T), NIHR BRC, and by Inserm, France. EPSRC (EP/R03558X/1), British Lung Foundation, Cambridge NIHR BRC. MRC Metabolic Diseases Unit [MC_UU_00014/5]. SOR is supported by MRC RG90505 and NIHR BRC RG85378. A.P.C and D.Y are supported by the MRC Metabolic Diseases Unit [MC_UU_0014/1]. BI is supported by ERC Consolidator Grant (No. 819775). EO is supported by MCIN/AEI Ramon y Cajal program (RYC2020-028884-I). RVG is supported by ISCIII P-FIS fellowship (FI17/00045). We thank Pfizer for the GDF15 and isotype control monoclonal antibody. Danna Breen and Bei Betty Zhang for advice and critical reading of the manuscript, Stephanie Joaquim, Matthew Lambert, Tao He and Laura Lin for design and preparation of antibodies. Meritxell Nus for advice and technical help. NIHR Core Biochemical Assay Laboratory, Addenbrooke’s Hospital, Cambridge, UK. BIOSTAT-CHF was funded by the European Commission (FP7-242209-BIOSTAT-CHF; and EudraCT 2010-020808-29). CNIC is supported by ISCIII, MCIN and Pro CNIC Foundation, and is a Severo Ochoa Center of Excellence (CEX2020-001041-S). Additional funding for measurement of GDF15 concentrations provided by Roche Diagnostics.

## Supplementary Figure legends

**Figure S1.**
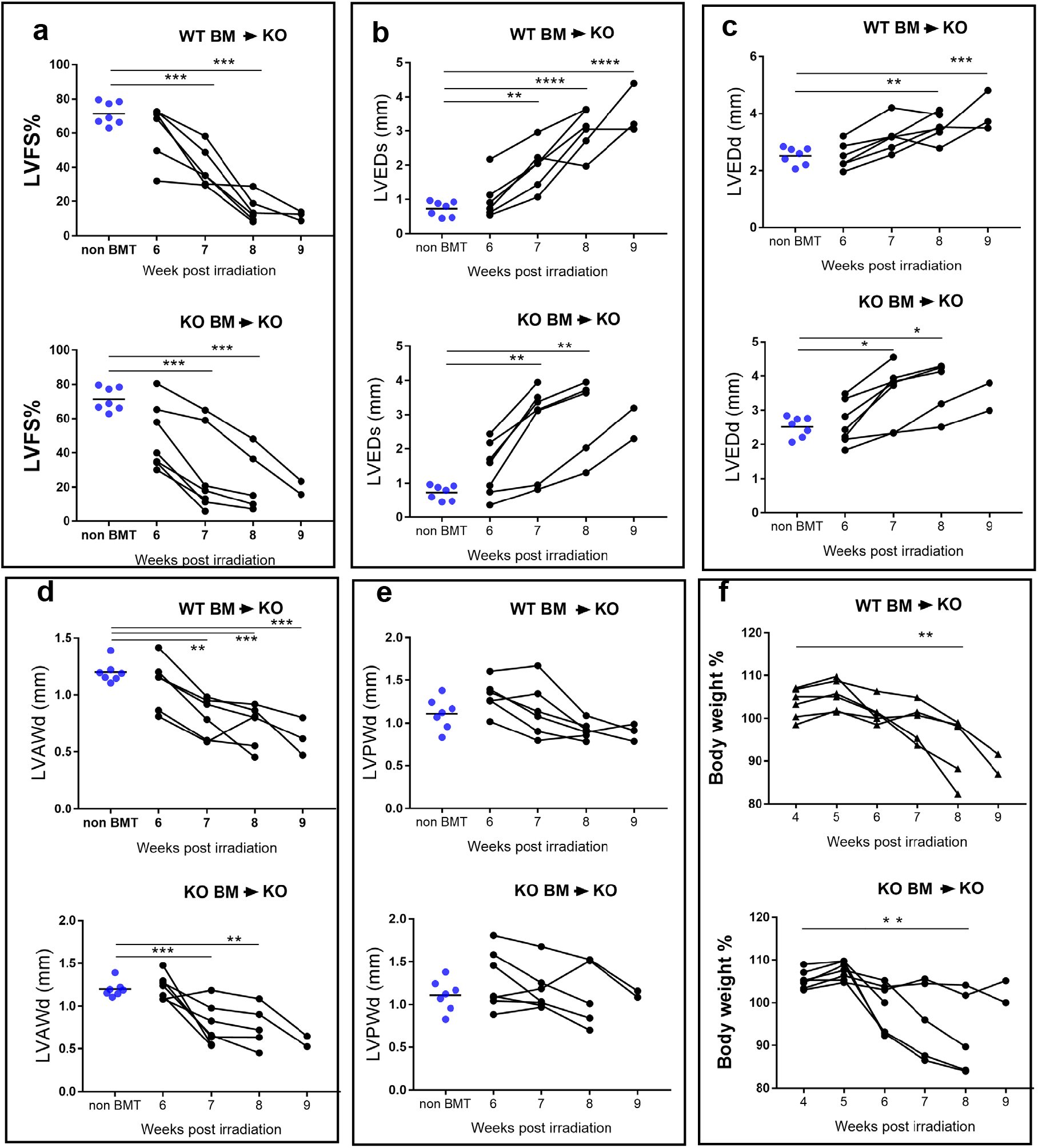
Related to Figure 1. Mice lacking functional PPP1R15A exhibit changes in heart function and geometry following whole body irradiation. *Ppp1r5a*^*ΔC/ΔC*^ male mice were irradiated (11Gy) followed by BM transfer using *Ppp1r15a*^+/+^ (WT BM) or *Ppp1r15a4*^*ΔC/ΔC*^ (KO BM) and monitored for (**a**) body weight, calculated as % of weight on day of irradiation and (**b**-**f**) heart function and geometry by echocardiography of parasternal short axis of the heart to assess (**b**) left ventricular fractional shortening, LVFS%. (**c**) LV end diameter at systole, LVEDs, (**d**) LV end diameter at diastole LVEDd, (E) LV anterior wall thickness, LVAWd, and (**f**) LV posterior wall diameter, LVPWd. Echocardiography-derived parameters over time were compared with parameters from non-irradiated *Ppp1r5a*^*ΔC/ΔC*^ male mice using one way ANOVA and Dunnett’s post comparison test. Body weight at each time point compared to weight at 4 weeks post-irradiation using one way ANOVA and Dunnett’s post comparison test. *p < 0.05 **p < 0.01, ***p < 0.001, ****p < 0.0001.

**Figure S2.**
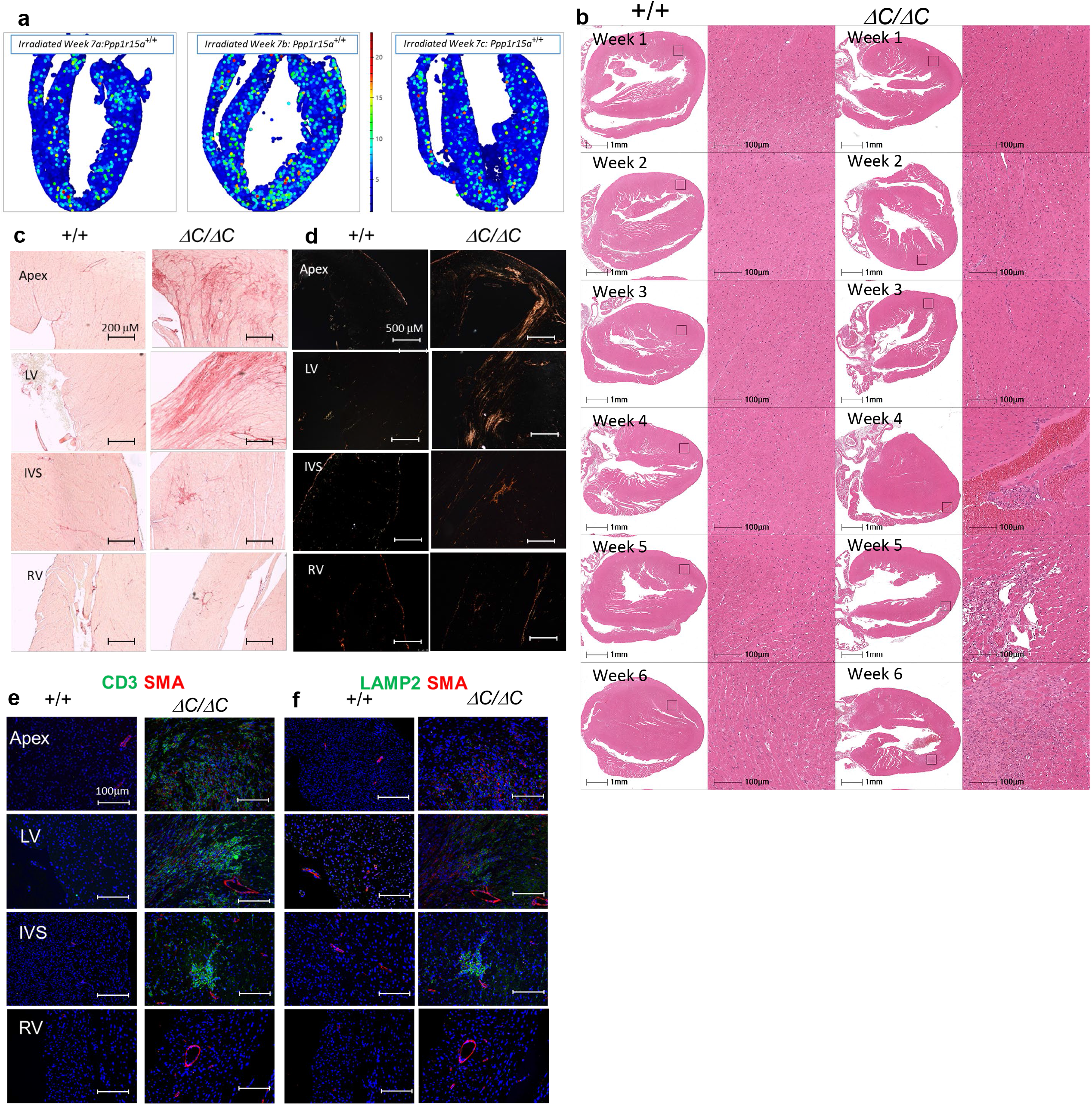
Related to Figure 2. Irradiated *Ppp1r15* a^+/+^ and *Ppp1r15a*^*ΔC/ΔC*^ mice exhibit histological differences in heart tissue. (**a**) Spatial plots of *Ppp1r15a* transcript levels assessed by Single Molecule In Situ Hybridisation (SM-ISH) in heart tissue sections derived from irradiated female WT mice, at 7 weeks post irradiation. (***b***) Comparison of single *Ppp1r15a*^*ΔC/ΔC*^ or *Ppp1r15a*^+/+^mice hearts from female mice, 1-6 weeks post-irradiation stained using H&E stain. Comparison of identical hearts shown in Figure 1c, 7 weeks post-irradiation, at 4 different locations – the apex, left ventricle (LV), interventricular septum (IVS) and right ventricle (RV). Sirius Red stain under (**c**) brightfield and (**d**) polarized light. Immunofluorescence staining for (**e**) CD3 (green) and (**f**) LAMP2 (Green), with smooth muscle actin (SMA) (red). The images were captured at equivalent locations and are representative of three hearts of each genotype.

**Figure S3.**
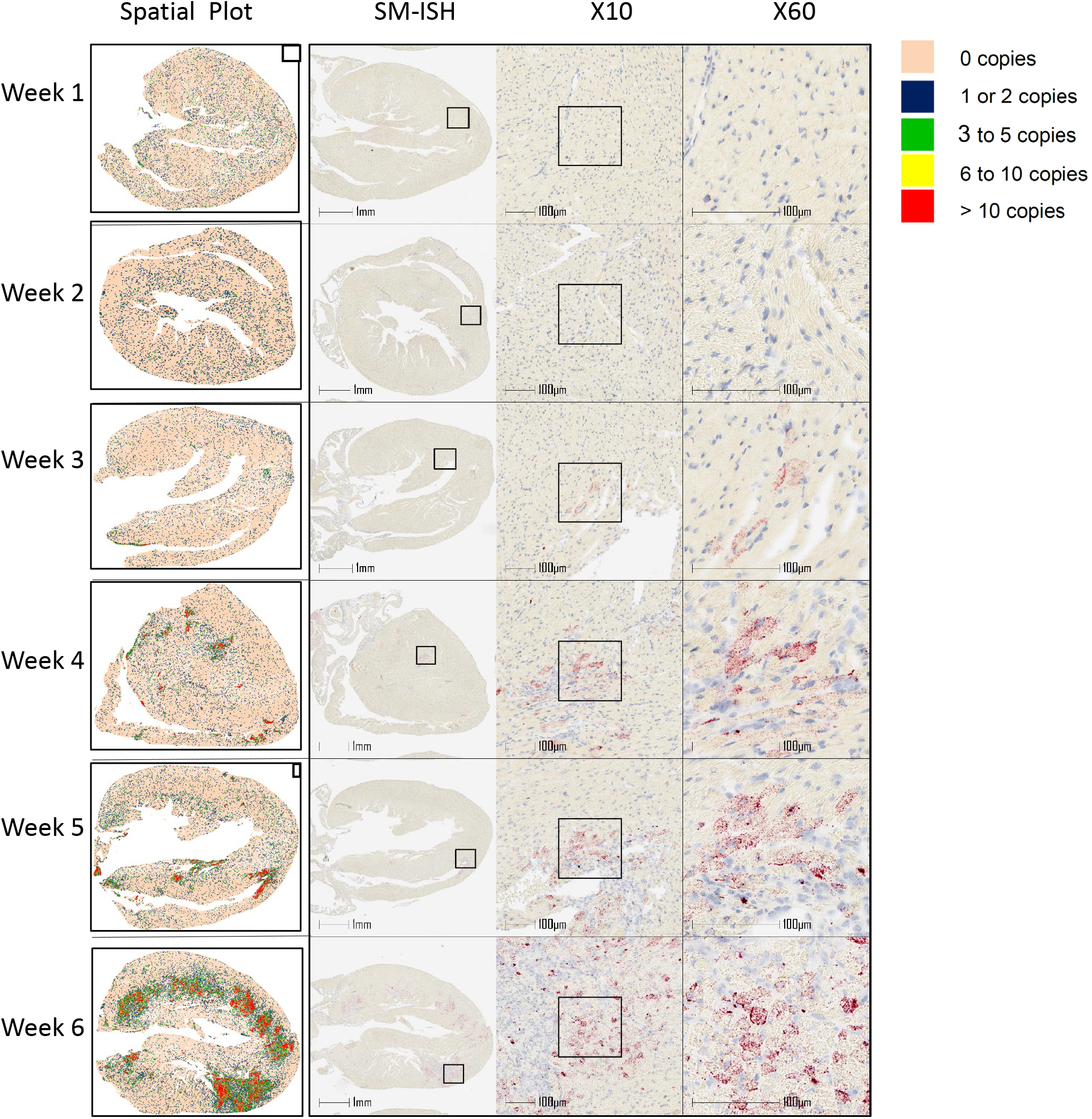
Related to Figure 3. Progression of *Gdf15* transcript expression in heart tissue following whole body irradiation of *Ppp1r15a*^*ΔC/ΔC*^ Mice. *Ppp1r15a*^*ΔC/ΔC*^ female mice (littermates) were irradiated (11Gy), followed by BM transfer and culled at the time points indicated, heart tissue was analysed by Small Molecule In Situ Hybridisation (SM-ISH) for *Gdf15* transcripts (red spots). Spatial plot shows distribution of quantified *Gdf15* mRNA transcripts, categorised as 0 copies, 1 to 2 copies, 3 to 5 copies, 6 to 10 copies and 10+ copies (see colour legend).

**Figure S4.**
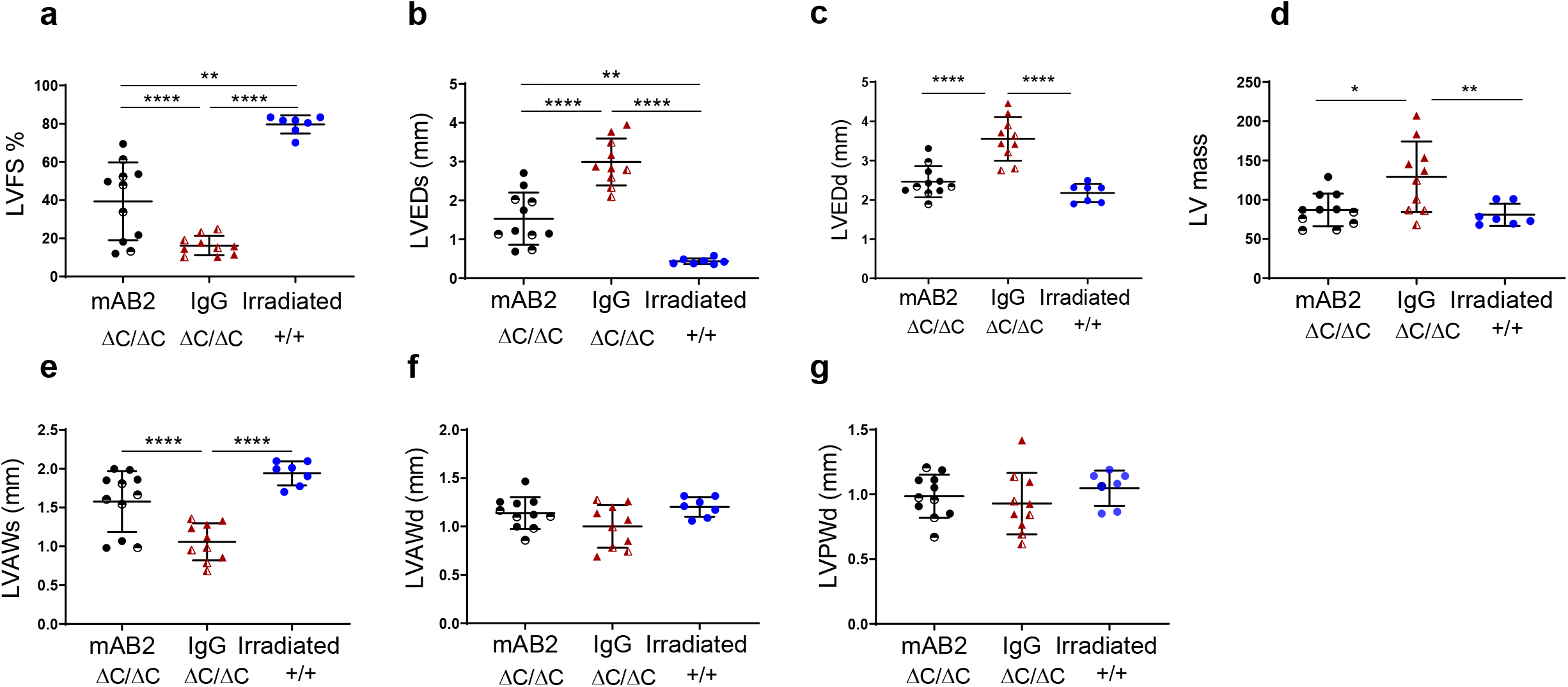
Related to figure 5. The GDF15 neutralising antibody, mAB2 changes parameters of heart function in irradiated mice lacking functional PPP1R15A. *Ppp1r15a*^*ΔC/ΔC*^ (^*ΔC/ΔC*^) or *Ppp1r15a*^+/+^ (*+/+*) mice were irradiated (11Gy) and reconstituted with *Ppp1r15a*^*+/+*^ bone marrow. At 4 weeks *Ppp1r15a*^*ΔC/ΔC*^ mice were given antibody that blocks GDF15 activity (mAB2) or an isotype control antibody (IgG) as described in Figure 1. Left ventricular heart function was assessed by echocardiography and expressed as (a) LVFS %, (b) LVEDs, (c) LVEDd, (d) LV mass, (e) LVAWs, (f) LVAWd and (g) LVPWd. Female *Ppp1r15a*^*ΔC/ΔC*^ mice are indicated using half-filled symbols. Data analysed using one way ANOVA and Tukey test post comparison analysis. All data show mean +/- SD. *p < 0.05 **p < 0.01, ***p < 0.001, ****p < 0.0001.

**Figure S5.**
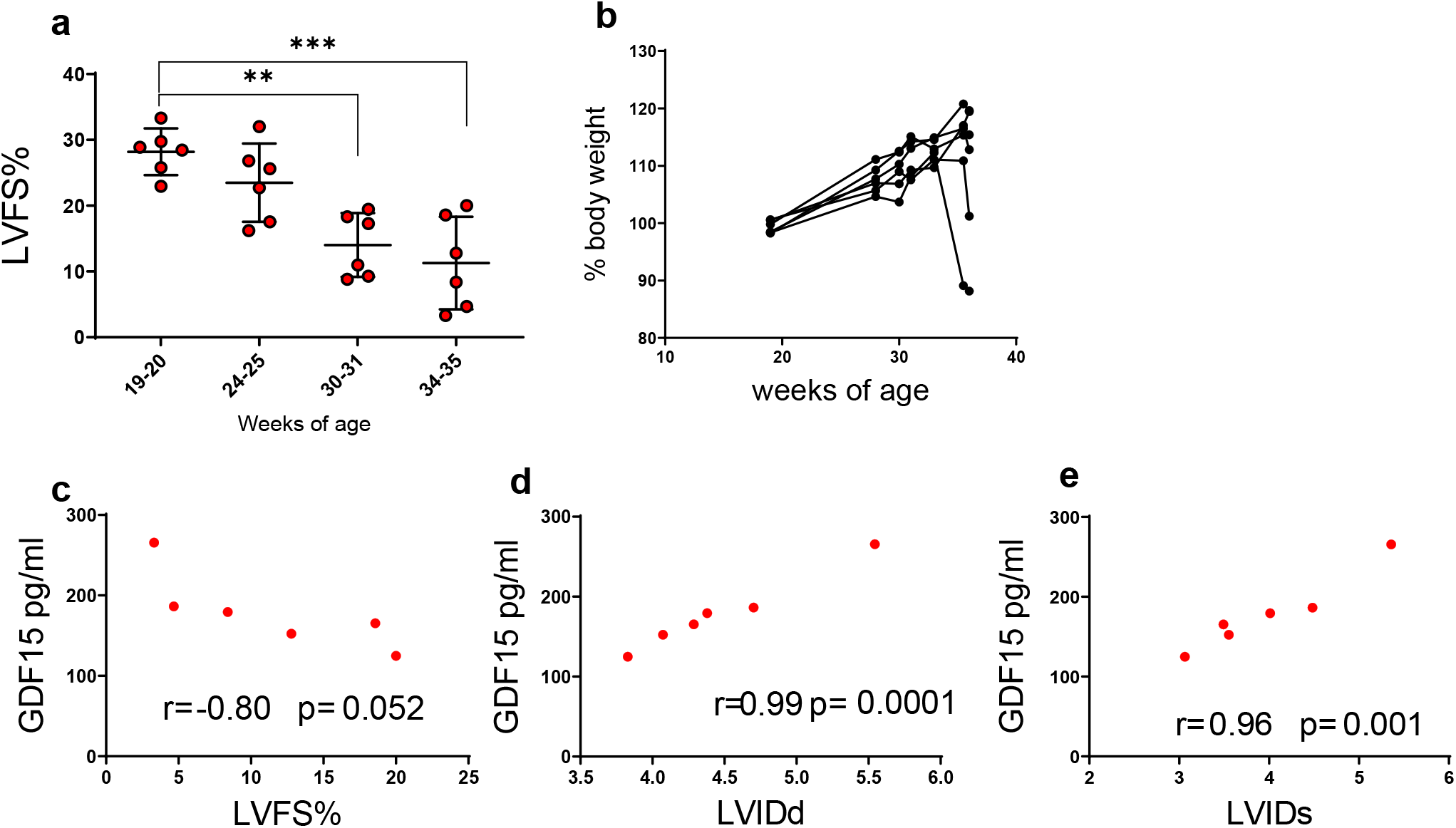
related to figure 6. GDF15 expression correlates with parameters of heart function and geometry in cYKO mice that have a cardiomyocyte specific deletion of Yme1l. cYKO mice that lack *Yme1l* in cardiomyocytes exhibit reduced heart function at 34-35 weeks of age (a). A proportion of the mice also exhibit dramatic weight loss (b). Plasma GDF15 collected at 34-35 weeks of age correlates with changes in parameters of heart geometry and heart function (c-e).

### Key resources table

**Table 1.**
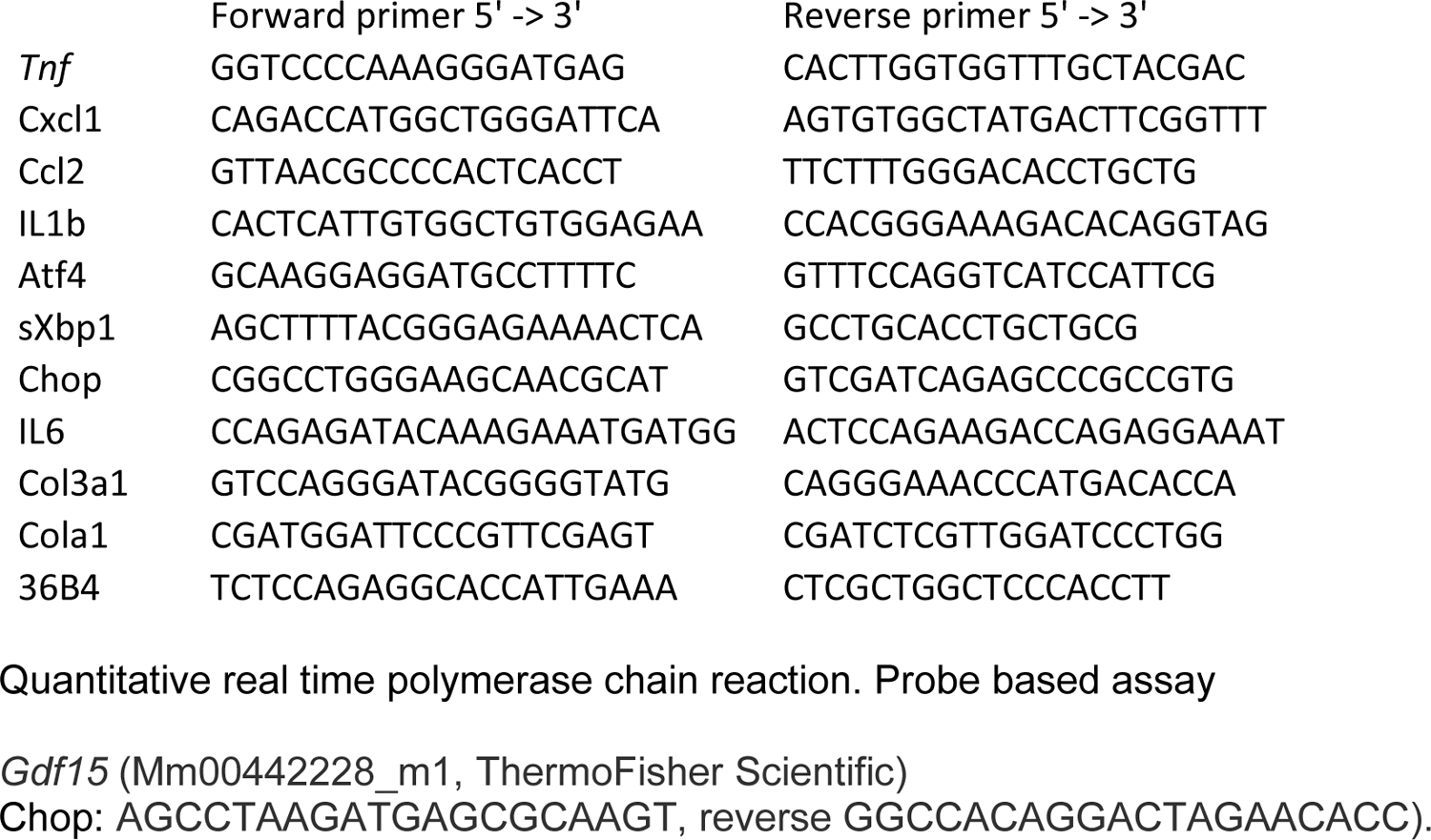
Quantitative real time polymerase chain reaction SYBR Green qPCR

